# Alpha-synuclein overexpression without vocalization deficits in a mouse model of parkinsonism

**DOI:** 10.1101/2025.06.17.660161

**Authors:** Brooke Rodgers, Allison Schaser

**Affiliations:** Department of Speech, Language, and Hearing Sciences, Purdue University, West Lafayette, Indiana, United States of America

**Keywords:** voice, ultrasonic vocalizations, parkinsonism, mouse model, alpha-synuclein

## Abstract

Voice deficits are common in Parkinson’s disease (PD) and significantly impact quality of life by increasing stress, social isolation, and caregiver burden. However, despite this impact, there are currently no treatments that target the underlying pathophysiology of PD in the vocalization system. The goal of this study was to examine the effect of one possible underlying mechanism responsible for the complex voice deficits that exist in PD; overexpression of the protein alpha-synuclein. Results show that overexpression of alpha-synuclein, prior to the development of alpha-synuclein aggregate pathology, does not result in significant vocalization deficits. A small but statistically significant increase in the total number of complex vocalizations was found in mice overexpressing alpha-synuclein compared to wildtype mice, but there were no differences in complexity ratio or any of the other specific vocalization parameters tested. Results provide a critical foundational understanding of the impact of overexpression versus aggregation of alpha-synuclein on voice deficits in PD. Future work will focus on manipulation of alpha-synuclein aggregate pathology, and not overexpression alone, to reduce or eliminate the burden of PD specific voice disorders.

**Summary Statement:** This study shows that overexpression of alpha-synuclein alone does not result in significant vocalization deficits, indicating that alpha-synuclein aggregate pathology within the vocalization system is required to induce vocalization deficits.

## Introduction

Voice deficits commonly occur in Parkinson’s disease (PD) and lead to significant impairments in quality of life (Ho *et al*., 1998; Miller, 2017). The complexity and variability of PD specific voice deficits make them inherently difficult to treat, which is highlighted by the controversy surrounding the inconsistent response of voice deficits to dopamine replacement therapy (Pinto *et al*., 2004; Ho, Bradshaw and Iansek, 2008).

Dopamine replacement therapy on its own is likely not enough to treat voice deficits because PD encompasses alpha-synuclein pathology at multiple levels and in multiple systems that could influence voice and communication, including extra-striatal areas in the brainstem and cortico-bulbar regions (Seidel *et al*., 2015; Postuma and Berg, 2016). To develop more robust and effective treatments for voice and communication deficits in PD, it is imperative that we understand the mechanisms underlying vocal deficits in PD. The inability of dopamine replacement therapy to effectively treat vocalization deficits leads to the hypothesis that some aspects of the vocal deficits in PD are not a result of the hallmark pathology in the striatal dopaminergic system, but instead result from abnormal alpha-synuclein in specific extra-striatal vocal communication areas.

Consistent with this hypothesis, extra-striatal alpha-synuclein pathology has been shown in cranial nerves and brainstem nuclei that control voice in both human patients (Mu *et al*., 2012) and in mouse models of alpha-synuclein overexpression (Grant *et al*., 2014). However, it is still unclear whether alpha-synuclein overexpression alone, without the presence of aggregate pathology, results in vocalization deficits.

The purpose of this study was to directly examine the independent role of alpha-synuclein overexpression on mouse ultrasonic vocalizations (USVs). To do this we characterized USVs in a previously established alpha-synuclein transgenic mouse model that overexpresses the aggressive disease-associated human A53T mutation via the mouse prion promotor and is tagged with a green fluorescent protein (GFP), known as the A53T SynGFP mouse model (Schaser *et al*., 2020). The A53T mutation is a point mutation in the *SCNA* gene at Alanine 53 that increases the probability of protein misfolding and is associated with familial PD (Giasson *et al*., 2002). While the A53T mutation leads to more rapid development of alpha-synuclein pathology after injection of pre-formed fibrils of alpha-synuclein, unlike other alpha-synuclein overexpression mouse models (Giasson *et al*., 2002; Chesselet *et al*., 2012; Sargent *et al*., 2017), the A53T SynGFP model does not develop spontaneous alpha-synuclein pathology over time (Schaser *et al*., 2020). As a result, we used the A53T SynGFP mouse model to determine if alpha-synuclein overexpression, prior to the onset of aggregate pathology, results in vocalization deficits.

## Results

USVs were elicited from a total of 29 transgenic and wildtype mice of both sexes using an adapted mating paradigm. For our purposes, only USVs in the 50 kHz range that were longer than 10 ms were included for analysis. USVs with only one acoustic element (flat, up/down, and U-shaped USVs) were considered simple vocalizations, and USVs with multiple acoustic elements (trailing, step up/down, and complex USVs) were considered complex vocalizations for analysis. The USV characteristics of interest were the total USV count, complexity ratio (calculated as the number of simple USVs divided by the number of complex USVs), duration (s), frequency range (kHz) (calculated as Fmax – Fmin), and the maximum intensity (dB). Over 19,000 vocalizations were identified by the USV identification software and included for statistical analysis. Of the vocalizations identified, 14,879 USVs were classified as simple and 4,245 USVs were classified as complex.

There were no statistically significant effects due to sex for any of the acoustic variables for either USV complexity type: total USV count (simple: *p* = 0.179; complex: *p* = 0.252), complexity ratio (*p* = 0.702), duration (simple: *p* = 0.314; complex: *p* = 0.943), frequency range (simple: *p* = 0.214; complex: *p* = 0.935), and maximum intensity (simple: *p* = 0.406; complex: *p* = 0.0524).

There were no statistically significant effects due to genotype for total USV count (simple: *p* = 0.279), complexity ratio (*p* = 0.392), duration (simple: *p* = 0.107; complex: *p* = 0.712), frequency range (simple: *p* = 0.167; complex: *p* = 0.816), and maximum intensity (simple: *p* = 0.57; complex: *p* = 0.194). There was a statistically significant effect due to genotype for the total number of complex vocalizations produced (*p* = 0.0429). Over the five recording days, animals overexpressing alpha-synuclein (transgene+) produced nine more complex USVs (or <2 more complex USVs per day) than wildtype (WT) animals on average. A table of the mean values for the genotype groups and *p*-values are shown in Table 1 and plots of the genotype group averages are shown in Figure 1.

**Table 1.** Group averages and P-Values for each USV characteristic calculated for the two genotype groups (transgene+ and wildtype) and averaged across sex.

|  | Simple USVs |  |  | Complex USVs |  |  |
| --- | --- | --- | --- | --- | --- | --- |
|  | Transgene+ | Wildtype | *P*-Value | Transgene+ | Wildtype | *P*-Value |
| **Average Total Count** | 510 | 518 | 0.279 | 150 | 141 | 0.0429* |
| **Average Complexity Ratio (simple/complex)** | 3.98 | 3.84 | 0.392 |  |  |  |
| **Average Duration (s)** | 0.0137 | 0.0134 | 0.107 | 0.0412 | 0.0405 | 0.712 |
| **Average Frequency Range (kHz)** | 3.01 | 2.73 | 0.167 | 23.0 | 22.8 | 0.816 |
| **Average Maximum Intensity (dB)** | -34.8 | -35.6 | 0.57 | -15.1 | -13.3 | 0.194 |

\*Asterisks indicate statistically significant differences due to genotype ( $p < 0.05$ ).

**Figure 1.**
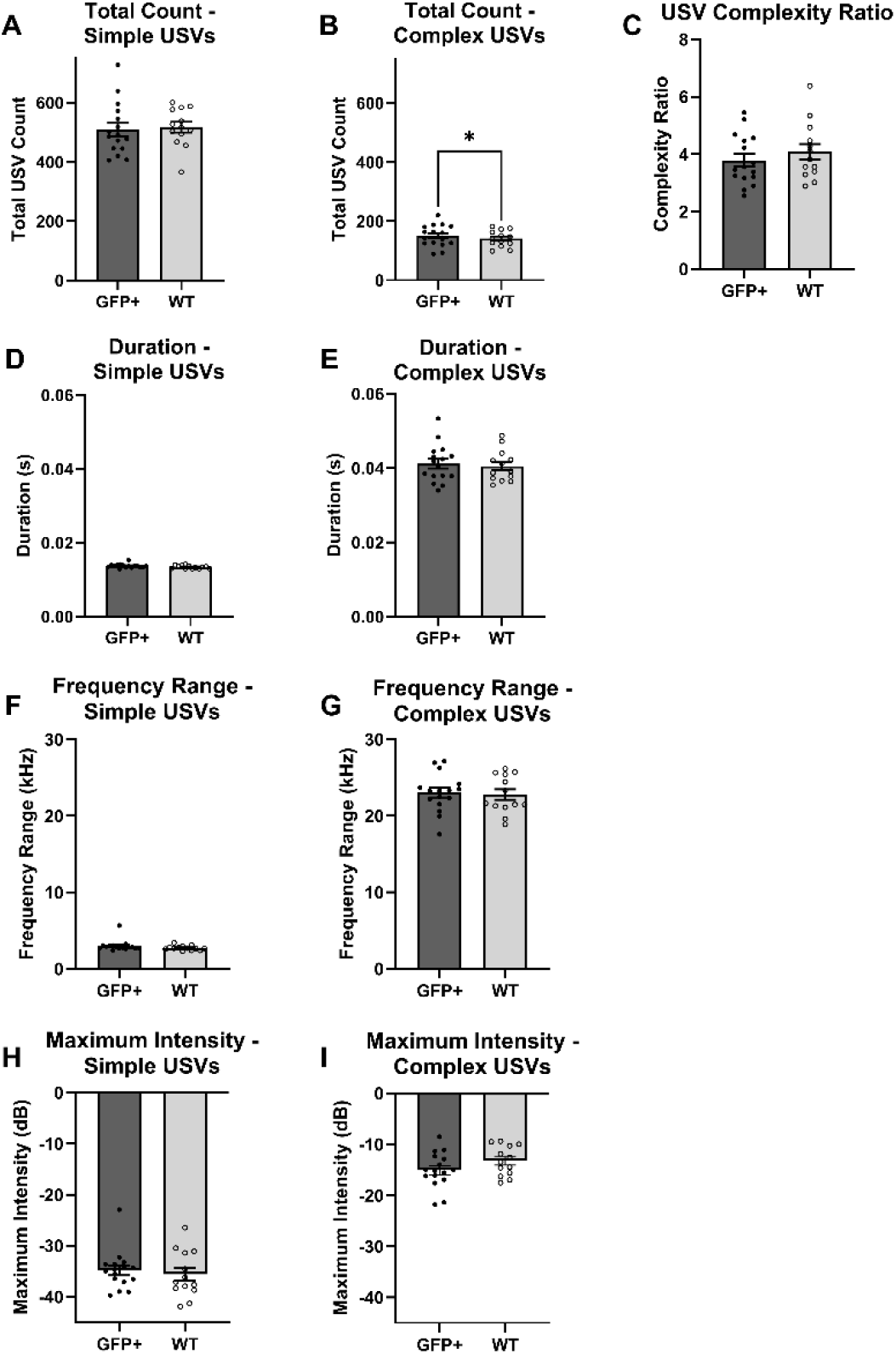
USV results averaged over each genotype group (GFP+ = transgene+ and WT = wildtype), pooled across sex. Results for simple (left panel) and complex (right panel) USVs were analyzed and plotted separately as total USV count (A, B), duration (D, E), frequency range (F, G), and maximum intensity (H, I). Group averages for USV complexity ratio are shown in C. Error bars represent the standard error of the mean value (s.e.m.) and points depict values for individual animals. The asterisk above the bars in plot B indicate a statistically significant difference between the group means (p < 0.05).

All animals produced more simple than complex USVs and had complexity ratios greater than 1. Across both groups, complex vocalizations had greater durations, frequency ranges, and maximum intensities than simple vocalizations, on average.

## Discussion

Results of this study show that overexpression of alpha-synuclein alone does not result in significant vocalization deficits. Using the A53T SynGFP mouse model, which overexpresses alpha-synuclein without developing spontaneous alpha-synuclein aggregate pathology, we were able to directly test the influence of alpha-synuclein overexpression alone on multiple USV parameters. We found that there was a small but statistically significant increase in the total number of complex vocalizations produced by the transgene+ mice compared to the WT mice in this study. However, the overall increase of 9 total complex USVs does not appear to represent a clinically meaningful change in vocalization and does not represent a vocalization deficit typically seen in clinical PD (Darley, Aronson and Brown, 1968; Ho *et al*., 1998; Plowman-Prine *et al*., 2009). All other USV parameters tested in the study were not different in A53T SynGFP mice compared to WT mice, indicating that overexpression of alpha-synuclein alone does not result in significant vocal deficits. This result is consistent with a previous study that examined vocalization in mice overexpressing alpha-synuclein (Grant *et al*., 2014). The Thy1-aSyn mice used in the previous study demonstrated early, progressive vocalization deficits compared to WT controls. However, even at the early time points tested in this study, vocalization deficits were linked to alpha-synuclein aggregation in the vocalization system, specifically in the periaqueductal gray (Grant *et al*., 2014).

Suggesting that overexpression alone was not sufficient to cause the vocalization deficits seen in this model but instead confirming the need for underlying alpha-synuclein aggregate pathology to cause vocalization deficits.

Future studies will benefit from the use of the A53T SynGFP mouse model tested here because it eliminates the concern that overexpression of alpha-synuclein alone influences USVs, but still allows for more rapid induction of alpha-synuclein pathology compared to WT mice (Schaser *et al*., 2020). Using this mouse model, we can control the location and timing of the development of alpha-synuclein aggregate pathology by taking advantage of additional methods to induce alpha-synuclein aggregate pathology. Alpha-synuclein is normally presynaptic, but in PD it forms somatic and neuritic inclusions, the hallmark lesions of this disease (Spillantini *et al*., 1997). Alpha-synuclein pathology propagation can be induced by the exogenous application of small, in vitro-generated alpha-synuclein pre-formed fibrils (PFFs). Previous research has shown that the application of PFFs causes aggregation of endogenous alpha-synuclein and resultant pathology first at the site of injection and then in connected brain regions in a time dependent manner (Luk *et al*., 2009; Sacino *et al*., 2014; Osterberg *et al*., 2015; Schaser *et al*., 2020). The specificity and inducibility of the A53T SynGFP mouse model and the PFF method make it an ideal system to examine alpha-synuclein aggregate pathology in striatal vs extra-striatal components of the vocal communication system.

The A53T SynGFP mouse model is ideal because it allows for rapid induction of pathology exclusively in one area while preserving the health of other areas. It also allows direct examination of the effect of alpha-synuclein pathology in discrete areas on vocal communication behavior, without the need to control for the additional effects of alpha-synuclein overexpression. Additionally, the GFP tag aids in genotyping mice and imaging alpha-synuclein pathology.

The results of the current study further support the hypothesis that vocalization deficits require the presence of alpha-synuclein pathology in specific areas of the vocalization system. This idea is supported by previous work in a rat PFF injection model which has shown that vocalization deficits do occur after PFF injection into the striatum, but these deficits only occurred once widespread pathology had developed over a 6 month period and did not appear to affect vocalization quality (bandwidth and intensity) (Paumier *et al*., 2015). This finding suggests a role for extra-striatal alpha-synuclein pathology in the overall scope of PD specific vocalization deficits.

Future studies should employ the current A53T SynGFP mouse model to determine the effects of alpha-synuclein pathology in discrete areas of the vocal communication system. Specifically, it will be important to induce alpha-synuclein pathology in areas along the vocal communication neuro-axis — including the vocal part of the periaqueductal gray and the layer V pyramidal neurons in the singing activated region of M1 (Arriaga, Zhou and Jarvis, 2012). Future studies will further elucidate the individual role of each discrete location on specific acoustic parameters in PD. The overall goal of this line of inquiry will be to determine the underlying cause of vocalization deficits in PD and to develop targeted treatments that address the variable and complex vocalization deficits that occur in PD.

The results of this study confirm the development of a mouse model that allows for manipulation of alpha-synuclein within discrete areas of the vocal communication system and enables the analysis of vocalizations in a relevant behavioral manner, without the need to control for the baseline level of alpha-synuclein overexpression. Our results provide a critical and necessary foundational understanding of the importance of aggregation and spread of alpha-synuclein pathology as one of the underlying mechanisms that leads to vocalization deficits in PD. Our data strongly suggest that the A53T SynGFP mouse model represents an ideal baseline system to test important fundamental questions related to how alpha-synuclein pathology propagation occurs within the vocal communication system and results in PD specific voice and communication disorders.

## Materials and Methods

### Mouse model

The procedures included in this study were performed in accordance with a protocol approved by the Oregon Health & Science University (OHSU) Institutional Animal Care and Use Committee. All experiments in this study were carried out at OHSU using a previously characterized alpha-synuclein transgenic mouse model, known as the A53T SynGFP mouse model (Schaser *et al*., 2020). This mouse model overexpresses the aggressive disease-associated human A53T mutation via the mouse prion promotor and is tagged with a green fluorescent protein (GFP). A total of 29 male and female mice, aged 3-6 months, were used in this study. Sixteen transgenic (transgene+) (7 male and 9 female) mice and 13 wildtype (WT) (7 male and 6 female) littermate control mice were used to determine if alpha-synuclein overexpression alone results in vocalization deficits.

### Ultrasonic Vocalization elicitation procedure

Ultrasonic vocalizations (USVs) were elicited using an adapted existing mating paradigm (Chabout, Jones-Macopson and Jarvis, 2017). Briefly, the procedure involved inducing a male or female mouse to vocalize by stimulating them with urine from a mouse of the opposite sex. Mouse urine was purchased and introduced into the testing cage via a cotton swab. Following interaction with the opposite sex, the scent elicited USVs from the mice. Five minutes of USVs were recorded from each animal over five consecutive days.

### Recording and USV identification

USVs were recorded using an ultrasonic recording system (Avisoft, Germany) with appropriate wide frequency response range (10 to 180 kHz) and the capability of producing spectrograms in real time. USVs were identified using the Sonotrack software package (Metris, Hoofddorp, Netherlands). After the final recording session, all recording files were processed automatically in Sonotrack using a custom set of parameters that identified USVs within a specified frequency range (25-100kHz) and above a threshold duration (10ms). The identified USVs were further classified by complexity based on acoustic parameters informed by previously published studies and the Metris categorization guidelines (Metris, 2017). USVs with only one acoustic component (flat, up/down, and chevron type vocalizations) were considered “simple” USVs, and USVs with multiple acoustic components (trailing, step up/down, split up/down, and complex type vocalizations) were considered “complex” USVs for our statistical analyses.

### Statistical analysis

The USV statistical analyses were performed using R Statistical Software (v4.3.0; R Core Team 2023, Vienna, Austria). After USV identification using Sonotrack, the output spreadsheets were concatenated to create a complete dataset including all USVs produced during each recording session. The variables of interest for statistical analysis included: 1) total USV counts, 2) the ratio of the number “simple” to “complex” USVs produced, 3) the duration of USVs in seconds (s), 4) the frequency range of USVs in kilohertz (kHz), and 5) the maximum intensity of USVs in decibels (dB). The total USV count was calculated by summing the number of USVs produced by each animal over the 5 recording days. The complexity ratio was calculated by dividing the number of simple USVs produced by the number of complex USVs produced by each animal each day (Ratio = # simple USVs / # complex USVs). The average complexity ratio was then calculated over the five recording days for each animal to use for statistical analysis. For the USV duration, frequency range, and maximum intensity, all the USVs produced were pooled over the five recording days to determine the average value for each animal. The average USV duration, frequency range, and maximum intensity were used for statistical analysis. Each variable was analyzed using a 2-way ANOVA model with the factors of sex (male or female) and genotype (wildtype or transgene+). Two-way interaction effects were included in the models. These models were used to evaluate any differences in the USVs produced by the mice due to genotype and/or sex at baseline. Group mean differences were considered statistically significant when *p* < 0.05.

Analyses were stratified between simple and complex USVs. Analyses were separated because of the acoustic parameters used to classify the USVs as simple or complex. Separate datasets were created to only include USVs classified as simple or complex, and the same statistical tests were performed on both datasets.

Estrous stage was not measured in the female mice during the recording period, but each stage of the 4–5-day cycles in the mice was captured over the five recording days (Zeng *et al*., 2023). We did not control for estrous stage in our statistical models, but instead averaged the acoustic variables over the five days to remove effects due to the estrous stage.

## Acknowledgments

The authors would like to acknowledge V. Unni and the animal care staff at OHSU for their assistance and support of the study. They would also like to acknowledge M. Norton and B. Hernandez for their assistance with data collection and analysis for this study.

## Competing Interests

No competing interests declared.

## Funding

This work was supported in part by the Collins Medical Trust Grants (AJS) and an American Speech-Language Hearing Foundation New Investigator Grant (AJS).

## Data and resource availability

All relevant USV data files, analysis script, and supporting images are publicly available through the Purdue University Research Repository (DOI: 10.4231/7V1E-7163; https://purr.purdue.edu/publications/4583/1).

## Notes

https://purr.purdue.edu/publications/4583/1

### Competing Interest Statement

The authors have declared no competing interest.

## References

Arriaga, G., Zhou, E.P. and Jarvis, E.D. (2012) ‘Of mice, birds, and men: the mouse ultrasonic song system has some features similar to humans and song-learning birds.’, PloS one, 7(10), p. e46610. Available at: 10.1371/journal.pone.0046610.

Chabout, J., Jones-Macopson, J. and Jarvis, E.D. (2017) ‘Eliciting and Analyzing Male Mouse Ultrasonic Vocalization (USV) Songs’, Journal of Visualized Experiments [Preprint], (123). Available at: 10.3791/54137.

Chesselet, M.-F., Richter, F., Zhu, C., Magen, I., Watson, M.B. and Subramaniam, S.R. (2012) ‘A Progressive Mouse Model of Parkinson’s Disease: The Thy1-aSyn (“Line 61”) Mice’, Neurotherapeutics, 9(2), pp. 297–314. Available at: 10.1007/s13311-012-0104-2.

Darley, F.L., Aronson, A.E. and Brown, J.R. (1968) ‘Motor speech signs in neurologic disease.’, The Medical clinics of North America, 52(4), pp. 835–44.

Giasson, B.I., Duda, J.E., Quinn, S.M., Zhang, B., Trojanowski, J.Q. and Lee, V.M.Y. (2002) ‘Neuronal alpha-synucleinopathy with severe movement disorder in mice expressing A53T human alpha-synuclein’, Neuron, 34(4), pp. 521–533. Available at: 10.1016/S0896-6273(02)00682-7.

Grant, L.M., Richter, F., Miller, J.E., White, S.A., Fox, C.M., Zhu, C., Chesselet, M.-F. and Ciucci, M.R. (2014) ‘Vocalization deficits in mice over-expressing alpha-synuclein, a model of pre-manifest Parkinson’s disease.’, Behavioral neuroscience, 128(2), pp. 110–21. Available at: 10.1037/a0035965.

Ho, A.K., Bradshaw, J.L. and Iansek, R. (2008) ‘For better or worse: The effect of levodopa on speech in Parkinson’s disease’, Movement Disorders, 23(4), pp. 574–580. Available at: 10.1002/mds.21899.

Ho, A.K., Iansek, R., Marigliani, C., Bradshaw, J.L. and Gates, S. (1998) ‘Speech impairment in a large sample of patients with Parkinson’s disease.’, Behavioural neurology, 11(3), pp. 131–137.

Luk, K.C., Song, C., O’Brien, P., Stieber, A., Branch, J.R., Brunden, K.R., Trojanowski, J.Q. and Lee, V.M.-Y. (2009) ‘Exogenous alpha-synuclein fibrils seed the formation of Lewy body-like intracellular inclusions in cultured cells.’, Proceedings of the National Academy of Sciences of the United States of America, 106(47), pp. 20051–6. Available at: 10.1073/pnas.0908005106.

Metris (2017) ‘Animal Behavior Analysis Solutions: Ultrasonic Vocalizations Automated Call Classification.’ Hoofddorp: Metris.

Miller, N. (2017) ‘Communication changes in Parkinson’s disease’, Practical Neurology, 17(4), pp. 266–274. Available at: 10.1136/practneurol-2017-001635.

Mu, L., Sobotka, S., Chen, J., Su, H., Sanders, I., Adler, C.H., Shill, H.A., Caviness, J.N., Samanta, J.E., Beach, T.G. and Arizona Parkinson’s Disease Consortium, the A.P.D. (2012) ‘Altered pharyngeal muscles in Parkinson disease.’, Journal of neuropathology and experimental neurology, 71(6), pp. 520–30. Available at: 10.1097/NEN.0b013e318258381b.

Osterberg, V.R., Spinelli, K.J., Weston, L.J., Luk, K.C., Woltjer, R.L. and Unni, V.K. (2015) ‘Progressive aggregation of alpha-synuclein and selective degeneration of lewy inclusion-bearing neurons in a mouse model of parkinsonism.’, Cell reports, 10(8), pp. 1252–60. Available at: 10.1016/j.celrep.2015.01.060.

Paumier, K.L., Luk, K.C., Manfredsson, F.P., Kanaan, N.M., Lipton, J.W., Collier, T.J., Steece-Collier, K., Kemp, C.J., Celano, S., Schulz, E., Sandoval, I.M., Fleming, S., Dirr, E., Polinski, N.K., Trojanowski, J.Q., Lee, V.M. and Sortwell, C.E. (2015) ‘Intrastriatal injection of pre-formed mouse α-synuclein fibrils into rats triggers α-synuclein pathology and bilateral nigrostriatal degeneration.’, Neurobiology of disease, 82, pp. 185–199. Available at: 10.1016/j.nbd.2015.06.003.

Pinto, S., Ozsancak, C., Tripoliti, E., Thobois, S., Limousin-Dowsey, P. and Auzou, P. (2004) ‘Treatments for dysarthria in Parkinson’s disease’, The Lancet Neurology, 3(9), pp. 547–556. Available at: 10.1016/S1474-4422(04)00854-3.

Plowman-Prine, E.K., Okun, M.S., Sapienza, C.M., Shrivastav, R., Fernandez, H.H., Foote, K.D., Ellis, C., Rodriguez, A.D., Burkhead, L.M. and Rosenbek, J.C. (2009) ‘Perceptual characteristics of Parkinsonian speech: A comparison of the pharmacological effects of levodopa across speech and non-speech motor systems’, NeuroRehabilitation, 24(2), pp. 131–144. Available at: 10.3233/NRE-2009-0462.

Postuma, R.B. and Berg, D. (2016) ‘Advances in markers of prodromal Parkinson disease’, Nature Reviews Neurology, 12(11), pp. 622–634. Available at: 10.1038/nrneurol.2016.152.

Sacino, A.N., Brooks, M., Thomas, M.A., McKinney, A.B., Lee, S., Regenhardt, R.W., McGarvey, N.H., Ayers, J.I., Notterpek, L., Borchelt, D.R., Golde, T.E. and Giasson, B.I. (2014) ‘Intramuscular injection of α-synuclein induces CNS α-synuclein pathology and a rapid-onset motor phenotype in transgenic mice.’, Proceedings of the National Academy of Sciences of the United States of America, 111(29), pp. 10732–7. Available at: 10.1073/pnas.1321785111.

Sargent, D., Verchère, J., Lazizzera, C., Gaillard, D., Lakhdar, L., Streichenberger, N., Morignat, E., Bétemps, D. and Baron, T. (2017) ‘“Prion-like” propagation of the synucleinopathy of M83 transgenic mice depends on the mouse genotype and type of inoculum’, Journal of Neurochemistry, 143(1), pp. 126–135. Available at: 10.1111/jnc.14139.

Schaser, A.J., Stackhouse, T.L., Weston, L.J., Kerstein, P.C., Osterberg, V.R., López, C.S., Dickson, D.W., Luk, K.C., Meshul, C.K., Woltjer, R.L. and Unni, V.K. (2020) ‘Trans-synaptic and retrograde axonal spread of Lewy pathology following pre-formed fibril injection in an in vivo A53T alpha-synuclein mouse model of synucleinopathy’, Acta neuropathologica communications, 8(1), p. 150. Available at: 10.1186/s40478-020-01026-0.

Seidel, K., Mahlke, J., Siswanto, S., Krüger, R., Heinsen, H., Auburger, G., Bouzrou, M., Grinberg, L.T., Wicht, H., Korf, H.-W., den Dunnen, W. and Rüb, U. (2015) ‘The Brainstem Pathologies of Parkinson’s Disease and Dementia with Lewy Bodies’, Brain Pathology, 25(2), pp. 121–135. Available at: 10.1111/bpa.12168.

Spillantini, M.G., Schmidt, M.L., Lee, V.M.-Y., Trojanowski, J.Q., Jakes, R. and Goedert, M. (1997) ‘|[alpha]|-Synuclein in Lewy bodies’, Nature, 388(6645), pp. 839–840. Available at: 10.1038/42166.

Zeng, P.-Y., Tsai, Y.-H., Lee, C.-L., Ma, Y.-K. and Kuo, T.-H. (2023) ‘Minimal influence of estrous cycle on studies of female mouse behaviors’, Frontiers in Molecular Neuroscience, 16, p. 1146109. Available at: 10.3389/fnmol.2023.1146109.

